# Validation of methods for Low-volume RNA-seq

**DOI:** 10.1101/006130

**Authors:** Peter A. Combs, Michael B. Eisen

## Abstract

Recently, a number of protocols extending RNA-sequencing to the single-cell regime have been published. However, we were concerned that the additional steps to deal with such minute quantities of input sample would introduce serious biases that would make analysis of the data using existing approaches invalid. In this study, we performed a critical evaluation of several of these low-volume RNA-seq protocols, and found that they performed slightly less well in metrics of interest to us than a more standard protocol, but with at least two orders of magnitude less sample required. We also explored a simple modification to one of these protocols that, for many samples, reduced the cost of library preparation to approximately $20/sample.

## 1 Introduction

Second-generation sequencing of RNA (RNA-seq) has proven to be a sensitive and increasingly inexpensive approach for a number of different experiments, including annotating genes in genomes, quantifying gene expression levels in a broad range of sample types, and determining differential expression between samples. As technology improves, transcriptome profiling has been able to be applied to smaller and smaller samples, allowing for more powerful assays to determine transcriptional output. For instance, our lab has used RNA-seq on single *Drosophila* embryos to measure zygotic gene activation [18] and medium-resolution spatial patterning [4]. Further improvements will allow an even broader array of potential experiments on samples that were previously too small.

For instance, over the past few years, a number of groups have published descriptions of protocols to perform RNA-seq on single cells (typically mammalian cells) [25, 23, 24, 11, 14]. A number of studies, both from the original authors of the single-cell RNA-seq protocols and from others, have assessed various aspects of these protocols, both individually and competitively [2, 27, 19]. One particularly powerful use of these approaches is to sequence individual cells in bulk tissues, revealing different states and cellular identies [3, 26].

However, we felt that published descriptions of single-cell and other low-volume protocols did not adequately address whether a change in concentration of a given RNA between two samples would result in a proportional change in the FPKM (or any other measure of transcriptional activity) between those samples. While there are biases inherent to any protocol, we were concerned that direct amplification of the mRNA would select for PCR compatible genes in difficult to predict, and potentially non-linear ways. For many of the published applications of single cell RNA-seq, this is not likely a critical flaw, since the clustering approaches used are moderately robust to quantitative changes. However, to measure spatial and temporal activation of genes across an embryo, it is important that the output is monotonic with respect to concentration, and ideally linear.

While it is possible to estimate absolute numbers of cellular RNAs from an RNAseq experiment, doing so requires spike-ins of known concentration and estimates of total cellular RNA content [21, 17]. However, many RNA-seq experiments do not do these controls, nor are such controls strictly necessary under reasonable, though often untested, assumptions of approximately constant RNA content. While ultimately absolute concentrations will be necessary to fully predict properties such as noise tolerance of the regulatory circuits [9, 8], many current modeling efforts rely only on scaled concentration measurements, often derived from *in situ*-hybridization experiments [7, 13, 12]. Given that, we felt it was not important that different protocols should necessarily agree on any particular expression value for a given gene, nor are we fully convinced that absolute expression of any particular gene can truly reliably be predicted in a particular experiment.

In order to convince ourselves that data generated from limiting samples would be suitable for our purposes, we evaluated several protocols for performing RNA-seq on extremely small samples. We also investigated a simple modification to one of the protocols that reduced sample preparation cost per library by more than 2-fold. Finally, we evaluated the effect of read depth on quality of the data. This study provides a single, consistent comparison of these diverse approaches, and shows that in fact all data from the low-volume protocols we examined are usable in similar contexts to the earlier bulk approach.

## 2 Results

### 2.1 Experiment 1: Evaluation of Illumina TruSeq

In our hands, the Illumina TruSeq protocol has performed extremely reliably with samples on the scale of 100ng of total RNA, the manufacturer recommended lower limit of the protocol. However, attempts to create libraries from much smaller samples yielded low complexity libraries, corresponding to as much as 30-fold PCR duplication of fragments. Anecdotally, less than 5% of libraries made with at least 90ng of total RNA yielded abnormally low concentrations, which we observed correlated with low complexity (Data not shown). To determine the lower limit of input needed to reliably produce libraries, we attempted to make libraries from 40, 50, 60, 70, and 80 ng of *Drosophila* total RNA, each in triplicate.

We considered the two libraries with lower than usual concentration to be failures. While a failure rate of approximately 1 in 3 might be acceptable for some purposes, we ultimately wanted to perform RNA sequencing on precious samples, where a failure in any one of a dozen or more libraries would necessitate regenerating all of the libraries. Furthermore, due to the low sample volumes involved (less than approximately 500pg of poly-adenylated mRNA), common laboratory equipment is not able to determine the particular point in the protocol where the failures occurred.

Thus, we consider 70 ng of total RNA to be the conservative lower limit to the protocol. While this is about 30% smaller than the manufacturer suggests, it is still several orders of magnitude larger than we needed it to be. We therefore considered using other small-volume and “single-cell” RNA-seq kits, which we had less experience with and less faith in the data.

### 2.2 Experiment 2: Competitive Comparison of Low-volume RNAseq protocols

We first sought to determine whether the low-volume RNAseq protocols available faithfully recapitulate linear changes in abundance of known inputs. We generated synthetic spike-ins by combining *D. melanogaster* and *D. virilis* total RNA in known, predefined proportions of 0, 5, 10, and 20% *D. virilis* RNA. For each of the low-volume protocols, we used 1ng of total RNA as input, whereas for the TruSeq protocol we used 100ng.

Although pre-defined mixes of spike-in controls have been developed and are commercially available [15], we felt it was important to ensure that a given protocol would function reproducibly with natural RNA, which almost certainly has a different distribution of 6-mers, which could conceivably affect random cDNA priming and other amplification effects. Furthermore, our spike-in sample more densely covers the approximately 10^5^ fold coverage typical of RNA abundances. It should be noted, however, that our sample is not directly comparable to any other standards, nor is the material of known strandedness. We assumed that the majority of each sample is from the standard annotated transcripts, but did not verify this prior to library construction and sequencing.

The different protocols had a variation in yield of libraries from between 6 fmole (approximately 3.6 trillion molecules) and 2,400 femtomoles, with the TruSeq a clear outlier at the high end of the range, and the other protocols all below 200 fmole (Table 2.2). All of these quantities are sufficient to generate hundreds of millions of reads—far more than is typically required for an RNA-seq experiment. We pooled the samples, attempting equimolar fractions in the final pool; however, due to a pooling error, we generated significantly more reads than intended for the TruSeq protocol, and correspondingly fewer in the other protocols. Unless otherwise noted, we therefore sub-sampled the mapped reads to the lowest number of mapped reads in any sample in order to provide a fair comparison between protocols.

We were interested in the fold-change of each *D. virilis* gene across the four samples, rather than the absolute abundance of any particular gene. Therefore, after mapping and gene quantification, we normalized the abundance *A*_*ij*_ of every gene *i* across the *j* = 4 samples by a weighted average of the quantity *Q*_*j*_ of *D. virilis* in sample *j*, as show in equation 1. Thus, within a given gene, a linear fit of *Â*_*ij*_ vs *Q*_*j*_ should have a slope of one and an intercept of zero.

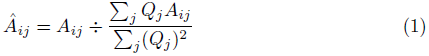

We filtered the *D. virilis* genes for those with at least 20 mapped fragments in the sample with 20% *D. virilis*, then calculated an independent linear regression for each of those genes. As expected, for every protocol, the mean slope was 1 (*t*-test, *p* < 5 × 10^−7^ for all protocols). Similarly, the average intercepts for all protocols was 0 (*t*-test, *p* < 5 × 10^−7^ for all protocols). Also unsurprisingly, the TruSeq protocol had a noticeably higher mean correlation coefficient (0.98 ± 0.02) than any of the other protocols (0.95 ± 0.06, 0.92 ± 0.09, and 0.95 ± 0.06 for Clontech, TotalScript, and SMART-seq2, respectively). The mean correlation coefficient was statistically and practically indistinguishable between the Clontech samples and the SMART-seq2 samples (*t*-test *p* = .11, Figure 2.2).

Indeed, the only major differentiator we could find between the low-volume protocols we measured was cost. For only a handful of libraries, the kit-based all inclusive model of the Clontech and TotalScript kits could be a significant benefit, allowing the purchase of only as much of the reagents as required. By contrast, the Smart-seq2 protocol requires the a la carte purchase of a number of reagents, some of which are not available or more expensive per unit for smaller quantities. Furthermore, there could potentially be a “hot dogs and buns” problem, where reagents are sold in non-integer multiples of each other, leading to leftovers. Many of these reagents are not single-purpose, however, so leftovers could in principle be repurposed in other experiments.

### 2.3 Experiment 3: Further modifications to the SMART-seq2 protocol

Although the SMART-seq2 was the cheapest of the protocols, we wondered whether it could be performed even more cheaply without compromising data quality. This would enable us to include more biological replicates in the future experiments for which we are evaluating these protocols. In the original protocol, we noticed that roughly 60% of the cost came from the Nextera XT reagents. Thus, reducing the cost of tagmentation was the obvious goal to target.

We made additional libraries, again starting with 1ng of total RNA. We amplified a single set of spike-in samples with 0, 5, 10, and 20% *D. virilis* total RNA as in experiment 2, and made a single an additional sample with 1% *D. virilis* RNA. Starting at the point in the SMART-seq2 protocol where tagmentation was started, we performed reactions in volumes 2.5× and 5× smaller, using proportionally less cDNA as well. Due to the low total yield, we increased the number of enrichment cycles from 6 to 8 (see methods).

When normalized to the same number of reads as in experiment 2, the protocols with diluted Nextera reagents performed effectively identically: for instance, the mean correlation coefficients were in both cases 0.96 ± 0.05 (Fig. 2 and Table 4). This is despite the additional cycles of enrichment, which improved yield.

**Figure 1.**
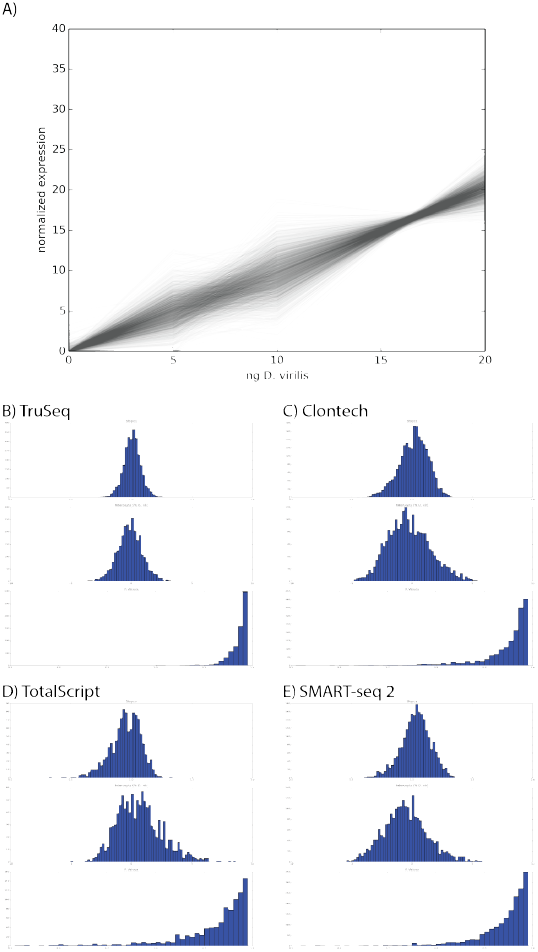
Comparison of linearity between different RNA-seq protocols. A) Normalized levels of gene expression *Â* across samples using the TruSeq protocol, where each line is for a different gene. B-E) Distributions of slopes, intercepts, and correlation coefficient for linear regressions of the data, as in panel A.

**Figure 2.**
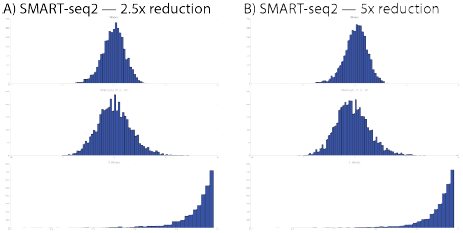
Distributions of slopes, intercepts, and correlation coefficients for experiment 3. Nextera XT reactions were reduced in volume by the indicated amount.

**Table 1:** Total TruSeq cDNA library yields made with a given amount of input total RNA. Yields measured by Nanodrop of cDNA libraries resuspended in 25*µL* of EB. The italicized samples were unusually low, and when analyzed with a Bioanalyzer, showed abnormal size distribution of cDNA fragments.

| Amount Input RNA | Replicate A | Replicate B | Replicate C |
| --- | --- | --- | --- |
| 40 ng | *57 ng* | 425 ng | 672 ng |
| 50 ng | 435 ng | 768 ng | 755 ng |
| 60 ng | *115 ng* | 663 ng | 668 ng |
| 70 ng | 300 ng | 593 ng | 653 ng |
| 80 ng | 468 ng | 550 ng | 840 ng |

**Table 2:**
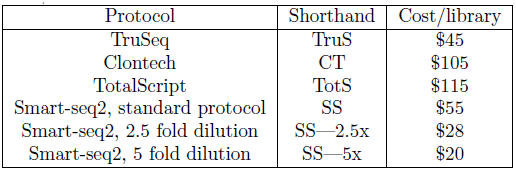
Summary of protocols used in experiments 2 and 3. Cost is estimated per sample assuming a large number of libraries at US catalog prices as of May 2014, and includes RNA extraction.

| Protocol | Shorthand | Cost/library |
| --- | --- | --- |
| TruSeq | TruS | \$45 |
| Clontech | CT | \$105 |
| TotalScript | TotS | \$115 |
| Smart-seq2, standard protocol | SS | \$55 |
| Smart-seq2, 2.5 fold dilution | SS—2.5x | \$28 |
| Smart-seq2, 5 fold dilution | SS—5x | \$20 |

**Table 3:** Sequencing summary statistics for samples. Protocols are the shorthands used in table 2. Reads indicates the total number of reads, and Mapped the total number of reads that mapped at least once to either genome. Experiments 2 and 3 were run in a single HiSeq lane each.

| Experiment | Protocol | % D. virilis | Yield (fmole) | Reads | Mapped |
| --- | --- | --- | --- | --- | --- |
| 2 | CT | 0 | 6.5 | 3,803,843 | 3,374,520 |
| 2 | " | 5 | 15.7 | 4,372,738 | 4,164,781 |
| 2 | " | 10 | 47.4 | 10,013,087 | 9,527,023 |
| 2 | " | 20 | 17.8 | 4,781,463 | 4,317,101 |
| 2 | TotS | 0 | 176.8 | 3,281,134 | 2,930,058 |
| 2 | " | 5 | 170.2 | 2,498,134 | 2,237,330 |
| 2 | " | 10 | 102.5 | 5,777,523 | 5,424,366 |
| 2 | " | 20 | 119.9 | 6,068,996 | 5,740,496 |
| 2 | TruS | 0 | 2,401.0 | 67,560,511 | 64,024,881 |
| 2 | " | 5 | 2,001.1 | 23,370,854 | 22,589,083 |
| 2 | " | 10 | 2,174.2 | 39,454,390 | 38,093,763 |
| 2 | " | 20 | 2,379.2 | 35,265,536 | 34,304,792 |
| 2 | SS2 | 0 | 34.3 | 2,439,518 | 2,297,087 |
| 2 | " | 5 | 59.6 | 2,550,023 | 2,419,889 |
| 2 | " | 10 | 67.9 | 2,534,628 | 2,444,568 |
| 2 | " | 20 | 39.8 | 2,504,340 | 2,389,850 |
| 3 | SS2—2.5x | 0 | 104.4 | 15,769,915 | 14,393,959 |
| 3 | " | 1 | 124.7 | 21,349,748 | 20,084,131 |
| 3 | " | 5 | 113.0 | 17,047,120 | 16,329,641 |
| 3 | " | 10 | 103.5 | 23,762,232 | 22,372,562 |
| 3 | " | 20 | 123.8 | 20,809,781 | 20,041,548 |
| 3 | SS2—5x | 0 | 59.4 | 19,214,155 | 17,324,598 |
| 3 | " | 1 | 58.6 | 23,832,274 | 22,364,220 |
| 3 | " | 5 | 65.4 | 18,149,452 | 17,157,450 |
| 3 | " | 10 | 28.8 | 15,821,419 | 14,869,864 |
| 3 | " | 20 | 57.2 | 22,466,345 | 21,620,603 |

**Table 4:**
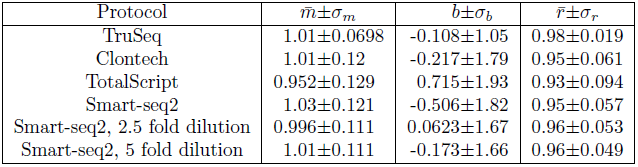
Distribution of fit parameters. A simple linear fit, 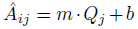 was computed for each gene *i*, and a correlation coefficent *r* calculated. For brevity, 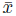 is the mean of some variable *x*, and *σ*_*x*_ is its standard deviation.

Because we used a common set of pre-amplified cDNA samples that was performed in a distinct pre-amplification from experiment 2, we can estimate the contribution of that pre-amplification to the overall variation. If, in fact, the pre-amplification is a major contributor to the variation, then we would expect to find that the correlation between, for instance, the slopes of two runs of the same experiment with different pre-amplifications would be significantly lower than the correlation between the slopes of two runs using the same pre-amplified cDNA pools.

Unsurprisingly, the sets of samples that used the same preamplification were more correlated with each other than with the set of samples that used a separate pre-amplification (Fig. 3). By analogy to dual-reporter expression studies[6], we term variation along the diagonal “extrinsic noise” (*η*_*ext*_ = std(*m*_1_ + *m*_2_)), and variation perpendicular to the diagonal “intrinsic noise” (*η*_*int*_ = std(*m*_1_ − *m*_2_)), being intrinsic to the pre-amplification step. Using that metric, the intrinsic noise is lower for the samples with the same pre-amplification (*η*_*int*_ = 0.09) than for the samples with different pre-amplifications (*η*_*int*_ = 0.16). Somewhat surprisingly, the extrinsic noise is higher for the samples with the same pre-amplification (*η*_*ext*_ = 0.20 vs *η*_*ext*_ = 0.16), perhaps due to the 2 additional cycles of PCR enrichment.

**Figure 3:**
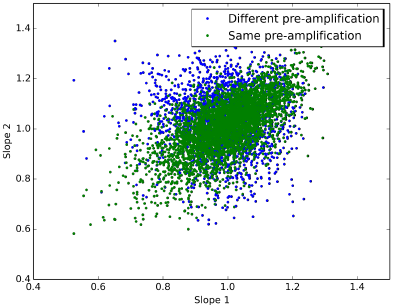
Estimating the source of preamplification noise. Plotted are the estimated slopes for each of the 3 samples. The 2.5× and 5× dilution samples used the same preamplified cDNA, but different tagmentation reactions, whereas the Full Size sample used different preamplification and tagmentation reactions.

## 3 Discussion

When sample size is not the limiting factor, it is clear that using well-established protocols that involve minimal sequence-specific manipulation of the sample yields the best results, both in terms of reproducibility and linearity of response. However, if it is not practical to collect such relatively large samples, we believe that any of the “single-cell” protocols we have tested should perform similarly, and can be used as a drop-in replacement. While preamplification steps do introduce some detectable variance, it is not vastly detrimental to the data quality, and does not introduce obvious sequence-specific biases.

Such methods should be strongly preferred if it is feasible to collect a suitably homogenous sample. While bulk tissues may be a mixture of multiple distinct cell types, this may or may not affect the particular research question an RNAseq experiment is designed to answer. In our hands, the lower limit of reliable library construction using the Illumina TruSeq kit is approximately 70 ng of total RNA; with non precious samples, the practical limit is likely to be even lower. Although we believe there is significant user-to-user variation, it seems unreasonable to expect order-of-magnitude improvements are possible in techniques for precious samples. We suggest that this limit may be related to cDNA binding to tubes or purification beads, but since the quantities are lower than the detection threshold of many standard quality control approaches, we cannot directly verify this, nor do we believe that knowing the precise cause is likely to suggest remediation techniques.

Compared to the regimes these protocols were designed for, we used a relatively large amount of input RNA—1 ng of total RNA—corresponding to approximately 50 nuclei of a mid-blastula transition *Drosophila* embryo. Previous studies have shown that this amount of RNA is well above the level where stochastic variation in the number of mRNAs per cell will strongly affect the measured expression of a vast majority of genes [19]. It is nevertheless a small enough quantity to be experimentally relevant. For instance, we have previously dissected single embryos into approximately 12 sections, yielding approximately 10 ng per section[4], and one could conceivably perform similar experiments on imaginal discs or antennal structures, which contain a similar amount of cells [16, 10].

One of the more striking results is that costs can be significantly reduced by simply performing smaller reactions, without noticeably degrading data quality. We do not suspect this will be true for arbitrarily small samples, such as from single cells. Instead, it is likely only true for samples near the high end of the effective range of the protocol. We have not explored where this result breaks down, and strongly caution others to verify this independently using small pilot experiments before scaling up.

## 4 Methods

### 4.1 RNA Extraction, Library Preparation, and Sequencing

We performed RNA extraction in TRIzol (Life Technologies, Grand Island, NY) according to manufacturer instructions, except with a higher concentration of glycogen as carrier (20 ng) and a higher relative volume of TRIzol to the expected material (1 mL, as in [18] and [4]). We quantified RNA concentrations using a fluorometric Qubit RNA HS assay (Life Technologies).

TruSeq libraries were prepared with the “TruSeq RNA Sample Preparation Kit v2” (Illumina Cat.#RS-122-2001) according to manufacturer instructions, except for the following modifications. All reactions were performed in half the volume of reagents. We find that this increases the effective concentration of RNA and cDNA. We performed all reactions and cleanups in 8-tube PCR strip tubes, which allowed us to reduce the volume of Resuspension Buffer to minimize volume left behind after each cleanup.

Clontech libraries were prepared with the “Low Input Library Prep Kit” (Clontech Cat.#634947). We generated cDNA by using TruSeq reagents until the cDNA synthesis step. Then, we used the Low Input Library Prep Kit to modify the cDNA into sequencing-competent libraries. We believe that a similar cDNA synthesis could be performed using oligo dT Dynabeads, RNA fragmentation reagents, and Superscript II (Life Technologies), for an approximate cost per sample of $15.

TotalScript libraries were prepared with the “TotalScript RNA-Seq Kit” and “TotalScript Index Kit” (Epicentre Cat.#TSRNA1296 and TSIDX12910). We followed the manufacturer’s instructions, and used the oligo dT priming option. We performed the mixed priming option in parallel, which yielded approximately 4-fold more library, but did not sequence them due to concerns of ribosomal contamination.

SMARTseq2 libraries were prepared according to the protocol in Picelli *et al.*(2014) [22]. Because we had already extracted and mixed the RNA, we began at step 5 with 3.7 *µL* of dNTPs and 1 *µL* of 37 *µM* oligo dT primer, yielding the same concentration of primer and oligo as originally reported. We used 18 cycles for the preamplification PCR in step 14, added 1ng of cDNA to the Nextera XT reactions in step 28, and used 6 and 8 cycles for the final enrichment in step 33 (experiments 2 and 3, respectively).

Libraries were quantified using a combination of Qubit High Sensitivity DNA (Life Technologies) and Bioanalyzer (Agilent Technologies, Sunnyvale, CA) readings, then pooled to equalize index concentration. Due to a pooling error in experiment 2, the TruSeq libraries were included at much higher abundance. Pooled libraries were then submitted to the Vincent Coates Genome Sequencing Laboratory for 50bp single-end sequencing according to standard protocols for the Illumina HiSeq 2500. Bases were called using HiSeq Control Software v1.8 and Real Time Analysis v2.8.

### 4.2 Mapping and Quantification

Reads were mapped using STAR [5] to a combination of the FlyBase reference genome version 5.54 for *D. melanogaster* and *D. virilis* [20]. We randomly sampled the mapped reads to use an equal number in each sample compared. We used HTSeq (command line options “`htseq-count --idattr=’gene\_name’ --stranded=no --sorted=pos`” to count absolute read abundance per gene [1].

## 5 Acknowledgements

## 6 Additional Information and Declarations

### 6.2 Author Contributions

Peter A. Combs conceived and designed the experiments, analyzed the data, and wrote the paper.

Michael B. Eisen conceived and designed the experiments and wrote the paper.

### 6.3 Data Deposition

